# pLM-DBPs: Enhanced DNA-Binding Protein Prediction in Plants Using Embeddings From Protein Language Models

**DOI:** 10.1101/2024.10.04.616755

**Authors:** Suresh Pokharel, Kepha Barasa, Pawel Pratyush, Dukka Kc

## Abstract

DNA-binding proteins (DBPs) play critical roles in gene regulation, development, and environmental response across various species, including plants, animals, and microorganisms. While various machine learning and deep learning models have been developed to distinguish DNA-binding proteins (DBPs) from non-DNA-binding proteins (NDBPs), most available tools have focused on human and mouse datasets. As a result, there are limited studies specifically addressing plant-based DNA-binding proteins, which restricts our understanding of their unique roles and functions in plant biology. Developing an efficient framework for improving DBP prediction in plants would enhance our knowledge and enable precise gene expression control, accelerate crop improvement, enhance stress resilience, and optimize metabolic engineering for agricultural advancement. In this work, we developed a tool that uses a protein language model (pLM) pre-trained on millions of sequences. We evaluated several leading models, including ProtT5, Ankh, and ESM-2, and leveraged their high-dimensional, information-rich representations to improve the accuracy of DNA-binding protein prediction in plants significantly. Our final model, pLM-DBPs, a feed-forward neural network classifier utilizing ProtT5-based representations, outperformed existing approaches with a Matthews Correlation Coefficient (MCC) of 83.8% on the independent test set. This represents a 10% improvement over the previous state-of-the-art model for plant-based DBP prediction, highlighting its superior performance compared to the existing approaches.

## Introduction

### Background

DNA-binding refers to the interaction between proteins and DNA, where specific proteins attach to particular sequences or structures within the DNA molecule that play a vital role in the structural composition of DNA. The proteins that can bind to DNA are called DNA-binding proteins [21, 6, 23]. In humans and other species, this interaction is crucial for various biological processes, including regulating gene expression, DNA replication, repair, and recombination. Understanding the nature of these proteins is crucial for insights into the regulation, the basis of many diseases, and the development of gene-based therapies [21, 27]. Similarly, in plants, DNA-binding proteins are essential for the regulation of nearly all aspects of plant life, from growth and development to environmental adaptation and defense responses [19]. Since experimental methods are often time-consuming and resource-intensive, numerous computational tools have been developed, with ongoing research focused on advancing these tools to enhance the efficiency and accuracy of their predictions.

Although there have been several computational tools developed to predict DBPs in Human and Mouse datasets, there are only a few tools to predict DBPs in plants. DNA-binding proteins (DBPs) in plants differ significantly compared to DBPs in other species [19]. Moreover, plant DBP prediction is crucial not only for basic biological understanding but also for practical applications in agriculture, environmental sustainability, and biodiversity conservation.

### Existing tools

Recent advancements in computational methods have significantly improved the prediction of DNA-binding proteins (DBPs) from sequences. Machine learning and deep learning-based tools have shown considerable promise in enhancing the accuracy of DBP prediction. However, many of these approaches rely on multiple handcrafted features, which require substantial time and effort in the data processing and model development process. For example, IDRBP-PPCT [25] uses a random forest model trained on combined features of Position-Specific Scoring Matrix (PSSM) and Position-Specific Frequency Matrix (PSFM) cross-transformation (PPCT) features. Similarly, StackDPPred [13] employs a combination of Position Specific Scoring Matrix (PSSM) features, Evolutionary Distance Transformation (EDT) features, Residue Probing Transformation (RPT) features, and a stacked ensemble of SVM, Logistic Regression, and KNN models to develop a DBP predictor. These methods incorporate a variety of machine learning techniques such as support vector machines (SVM), random forests (RF), adaptive boosting (ADB), and light gradient boosting (LGB).

iDRBP MMC [29] is a deep learning model that predicts both DNA and RNA binding proteins (RBPs) using a multi-label classification approach. It also uses PSSM features to train a convolutional neural network (CNN) for predicting four classes: DBPs, RBPs, non-DNA-binding proteins (NDBPs), and dual RNA-DNA binding proteins (DRBPs). Similarly, DeepDRBP-2L [28] is a CNN-biLSTM (Convolutional Neural Network-Bidirectional Long Short-Term Memory) model trained on PSSM features to predict these same classes. PDBP-Fusion [9] also combines CNN and biLSTM networks but uses one-hot encoded features for training. Recently, Wu et al. [27] proposed two approaches: hierarchical and multi-class CNN-LSTM model trained on PSSM features, achieving balanced prediction accuracy for both DBPs and RBPs within a single model.

Despite the development of numerous tools for DNA-binding protein (DBP) prediction, only a limited number are specifically tailored to plant DBPs [14, 19]. To our knowledge, PlDBPred [19] is the most recent tool developed specifically for plant DBP prediction which is a support vector machine model trained on the integrated PSSM features. Since existing methods predominantly rely on handcrafted features, including PSSM and physicochemical properties, the effective application of protein language model-based representations for this task, which have demonstrated significant success in various bioinformatics applications [17, 7, 18, 5], remains an area that needs further exploration. Our approach focuses on leveraging these sequence-based pLMs more efficiently to enhance plant DBP prediction.

### Proposed work

Latest developments in large language models have revolutionized Natural Language Processing, yielding impressive results across a wide range of tasks. This progress has also influenced the field of computational biology, specifically the development of protein language models (pLMs). These models are trained on a large corpus of protein sequences that have demonstrated an extraordinary ability to capture meaningful representations of protein sequences. Notable examples include Ankh[3], ESM versions-1/2/3 (Evolutionary Scale Modeling) [10], and ProtT5 [3], which have shown exceptional performance across various protein-related tasks, such as predicting post-translational modifications [17, 20, 16], binding residues in disordered regions [7], and protein structure [2]. Building on the success of these protein language models, this study explores their potential for predicting DNA-binding proteins in plants. We investigated several pLMs and developed a simple yet efficient model for plant DNA-binding protein prediction, named pLM-DBPs. Our results show that using pLM-based representations significantly improves prediction accuracy, suggesting that these models are adept at capturing hidden features within protein sequences, leading to more precise and reliable predictions.

## Materials and Methods

### Dataset

The training and testing sets were adopted from PlDBPred [19], which compiled its data from the UniProt database [1]. The dataset includes DNA-binding proteins (positive set) and non-DNA-binding proteins (negative set) derived from plants across 35 species. After pre-processing, the PlDBPred dataset contained 849 DNA-binding proteins and 1,848 non-DNA-binding proteins. The negative set was balanced by randomly selecting an equal number of non-DNA-binding proteins to match the number of DNA-binding proteins. An independent test set was also compiled from the same database, following identical processing steps. This independent dataset consisted of 997 sequences, including 500 non-DNA-binding and 497 DNA-binding proteins. Table 1 reports the distribution of DNA-binding proteins and non-DNA-binding proteins in train and test set.

**Table 1.** Number of plant-based DNA-binding and non-DNA binding proteins for training and test set used in this study.

| Split | # DBPs | # NDBPs |
| --- | --- | --- |
| Train | 849 | 849 |
| Test | 500 | 497 |
| Total | 1349 | 1346 |

### Proposed model workflow architecture

Figure 1 provides a high-level overview of the study’s workflow, divided into three main stages: data preparation and feature extraction, model development, and evaluation.

**Fig. 1.**
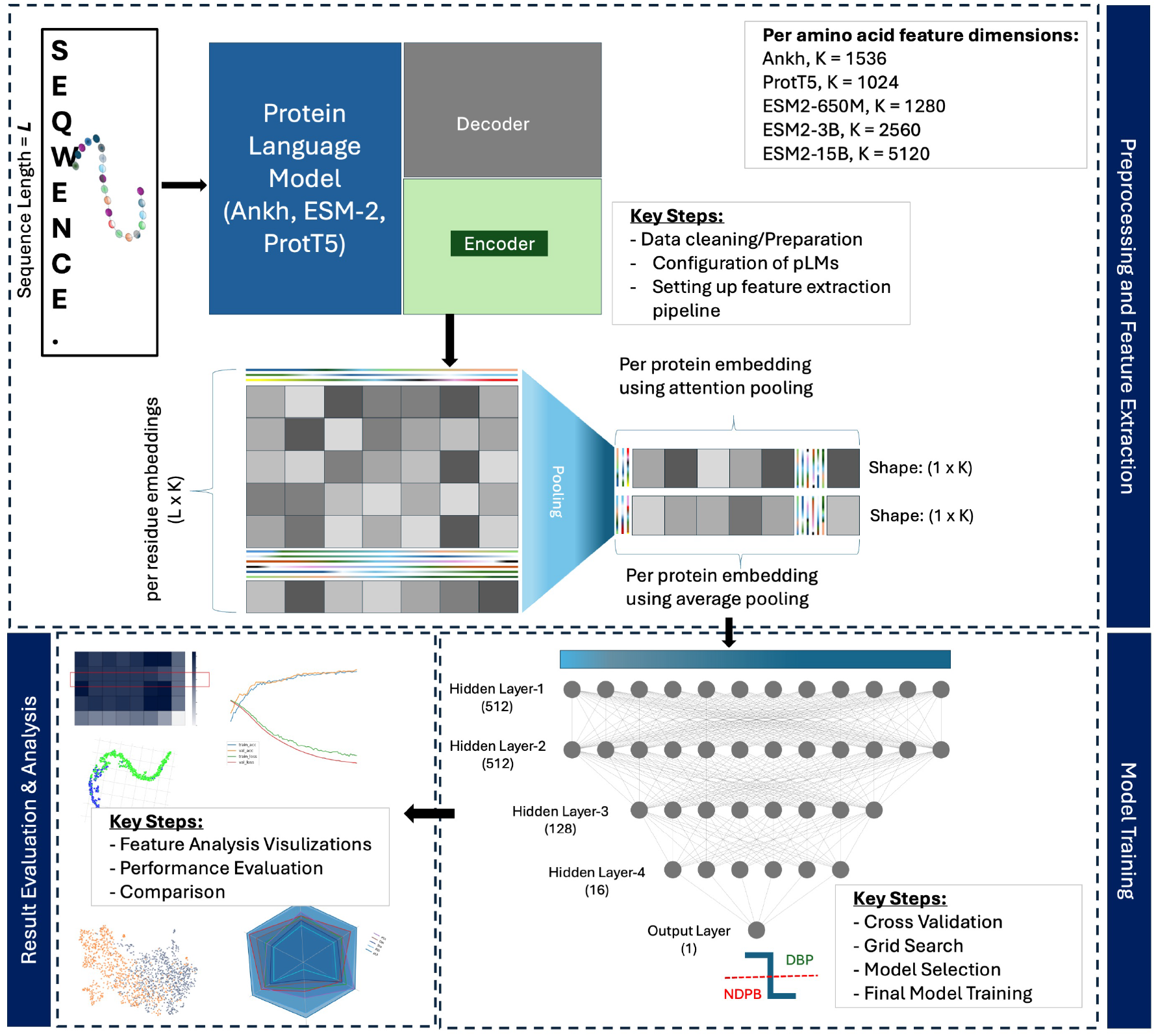
Overview of the study’s workflow illustrating key stages, including feature extraction using protein language models (Ankh, ESM-2, ProtT5), model development/training, evaluation

The figure illustrates the process starting with protein sequences, which are prepared as inputs for a protein language model (pLM). Next, the extracted residue-level representations from pLMs are pooled to create a protein-level representation. Then, this protein level representation is subsequently used as features to train and develop a downstream model for predicting DNA-binding proteins (DBPs), followed by a thorough performance evaluation of the model. The main components of Figure 1 are detailed in the subsequent subsections, along with the experimental methodologies for each stage.

#### Protein language models

In the rapidly advancing field of computational biology, the emergence of large protein language models has profoundly influenced research and applications across various tasks of bioinformatics. In this work, we utilize protein language models (pLMs) that have been primarily trained on a large corpus of unlabeled protein sequences. These models are capable of representing sequences uncovering hidden information patterns. In this work, we input embeddings from three non-fine-tuned pLMs: Ankh, ESM-2’s three versions, and ProtT5. Ankh is a general-purpose protein language model that contains a relatively smaller number of parameters compared to other competitive models [3]. It was trained on the Uniref50 dataset, which comprises 45 million protein sequences. This protein language model has two versions available: Ankh_base and Ankh_large. Ankh_base and Ankh_large versions have 450 million and 1.15 billion trainable parameters which output an amino acid residue to an embedding dimension of 768 and 1536 respectively. In this study, we used the Ankh_large version (referred to as Ankh in this paper). Evolutionary Scale Modeling (ESM) is a bidirectional encoder representation from a transformers architecture (BERT) based model also trained on the UniRef50 dataset. ESM models are designed to predict masked amino acids by considering the entire sequence context. In this work, we explored three variations of ESM2 models [11] including esm2_t33_650M_UR50D, esm2_t36_3B_UR50D, and esm2_t48_15B_UR50D. Hereafter, these models are referred to as ESM2-650M, ESM2-3B, and ESM2-15B in the context of this paper, respectively. ProtT5-XL-U50 [4] (referred as ProtT5 in this paper) is another language model pre-trained on the UniRef50 database, employing a masked language modeling (MLM) approach to predict hidden amino acids within sequences. Based on Google’s T5-3B architecture, ProtT5 retains key structural elements using 3 billion trainable parameters and 24 layers in both its encoder and decoder. It differs from the original T5 by using a BART-like MLM objective instead of span denoising. In this study, only the encoder component of pLMs was utilized for feature extraction. Detailed description of these models is summarized in Table 2.

**Table 2.** Overview of protein language models (pLMs) used in this study summarizing their architectures, number of parameters, layer counts, training datasets, embedding dimensions, and other details.

| pLM | Dataset | Architecture | Developer | # Parameters (Billions) | # Layers | Embedding Dimension per Residue |
| --- | --- | --- | --- | --- | --- | --- |
| Ankh-Base | UniRef50 | ConvBERT | Rostlab | $\approx 0.45\text{B}$ | - | 768 |
| Ankh-Large | | | | $\approx 1.2\text{B}$ | - | 1536 |
| ProtT5 | UniRef50 | T5-3B | Rostlab | $\approx 3\text{B}$ | 24 | 1024 |
| ESM2-650M | UniRef50 | RoBERTa | Meta | 0.65B | 33 | 1280 |
| ESM2-3B |  |  |  | 3B | 36 | 2560 |
| ESM2-15B |  |  |  | 15B | 48 | 5120 |

Ankh, ProtT5 models are taken from Hugging Face (https://huggingface.co)
ESM models are downloaded from their GitHub repository ( https://github.com/evolutionaryscale/esm)

### Feature extraction using protein language models

In this study, protein language models are used to extract the protein representation in the form of feature vectors for protein sequences from the hidden layers of their pre-trained encoder. Those representations serve as feature inputs to our downstream model for protein-DNA prediction in plants. Our proposed model, pLM-DBPs, utilizes pLMs to encode each amino acid in a protein sequence to corresponding feature vectors. As illustrated in Figure 1, a protein sequence of length *L* is fed to the pLM, resulting in vector embeddings of dimension *K* × 1 for each amino acid. Subsequently, these residue-level vector embeddings are consolidated into a protein-level representation using a pooling technique. We explored two types of pooling: average pooling and attention pooling. Both of these processes yield a single vector representation for the entire protein sequence, providing the full sequence representation. For example, in the case of ESM2-3B, each protein sequence is represented by a feature vector of dimension 2560 × 1. Since the results from different pooling techniques did not make a significant difference, we considered the average pooling mechanism for the experiments including our final model. Details about these pooling methods and the results of additional experiments explored using attention pooling are provided in supplementary information (Table T1).

### Model selection

#### Cross-validation and hyperparameter-tuning

In this study, a 5-fold cross-validation is performed to assess the performance of various machine learning models trained on our non-DBP and DBP datasets. The dataset was randomly split into five equal folds, with one fold serving as the validation set in each iteration, while the remaining four folds form the training set. The performance on each validation set is presented in Table 4, with final results representing the averages across all folds. To select the best model and hyperparameters, we performed a grid search training over a wide range of configurations for support vector machines (SVM), random forest (RF), and feed-forward artificial neural network (ANN) models. We used KerasTuner [15] to optimize the hyperparameters of the neural network. The best-performing model was selected based on the highest accuracy from the cross-validation, and its optimal hyperparameters were used to develop the final model.

#### Final model training

Protein level features extracted from protein language models are further trained to make classifiers as discussed in Section 2.5. Based on the various model selection results, we developed an Artificial Neural Network model trained using ProtT5 features that classifies the DBPs from NDBPs. Our final model consists of four hidden layers with 512, 512, 128, and 16 neurons, in addition to the input and output layers. The dropout layers were introduced in between layers to prevent the model from model overfitting. The details of the model summary are provided in the supplementary information Table T2. The training accuracy/loss curves are provided in supplementary information in Figure S1.

#### Model evaluations

In this study, we used a range of evaluation metrics to assess the performance of our models, consistent with established methods in the field. Metrics such as Sensitivity (Sn), Specificity (Sp), Matthew’s Correlation Coefficient (MCC), and Accuracy (ACC) are detailed in Table 3 with their respective formulas, providing a comprehensive evaluation of model performance. Using standard classification terminology, TP (True Positive) refers to correctly predicted DNA-binding sequences, FP (False Positive) to incorrectly predicted DNA-binding sequences, TN (True Negative) to correctly identified non-DNA-binding sequences, and FN (False Negative) to misclassified non-DNA-binding sequences.

**Table 3.** Performance metrics and their formula used for model selection and model evaluation used in this study.

| Metrics | Formula |
| --- | --- |
| Sensitivity | $\frac{TP}{TP+FN}$ |
| Specificity | $\frac{TN}{TN+FP}$ |
| Accuracy | $\frac{TP+TN}{TP+FP+TN+FN}$ |
| Precision | $\frac{TP}{TP+FP}$ |
| F-Score | $\frac{2 \cdot \text{Precision} \cdot \text{Recall}}{\text{Precision} + \text{Recall}}$ |
| MCC | $\frac{(TP \cdot TN) - (FP \cdot FN)}{\sqrt{(TP+FP)(TP+FN)(TN+FP)(TN+FN)}}$ |

## Results and Discussion

### 5-fold cross-validation results

The five-fold cross-validation results for hyperparameter optimization and model selection learning models, including SVM, RF, and ANN, were trained with a wide range of hyperparameters to evaluate the effectiveness of different feature types. Since all the machine learning algorithms demonstrated consistently strong results with minimal variation, we opted not to train more complex architectures. This allowed us to focus on simpler, well-performing models without the need for additional complexity.

In Table 4, while multiple models demonstrate comparable results, the ANN model trained on ProtT5 features stands out with the highest AUROC and AUPR and competitive performance in other metrics. As noted by some previous works [7, 24, 26], the rich information content from protein language model-based embeddings allowed the downstream model to be relatively simple architecture, allowing for shallower models are effective enough downstream tasks.

**Table 4.**
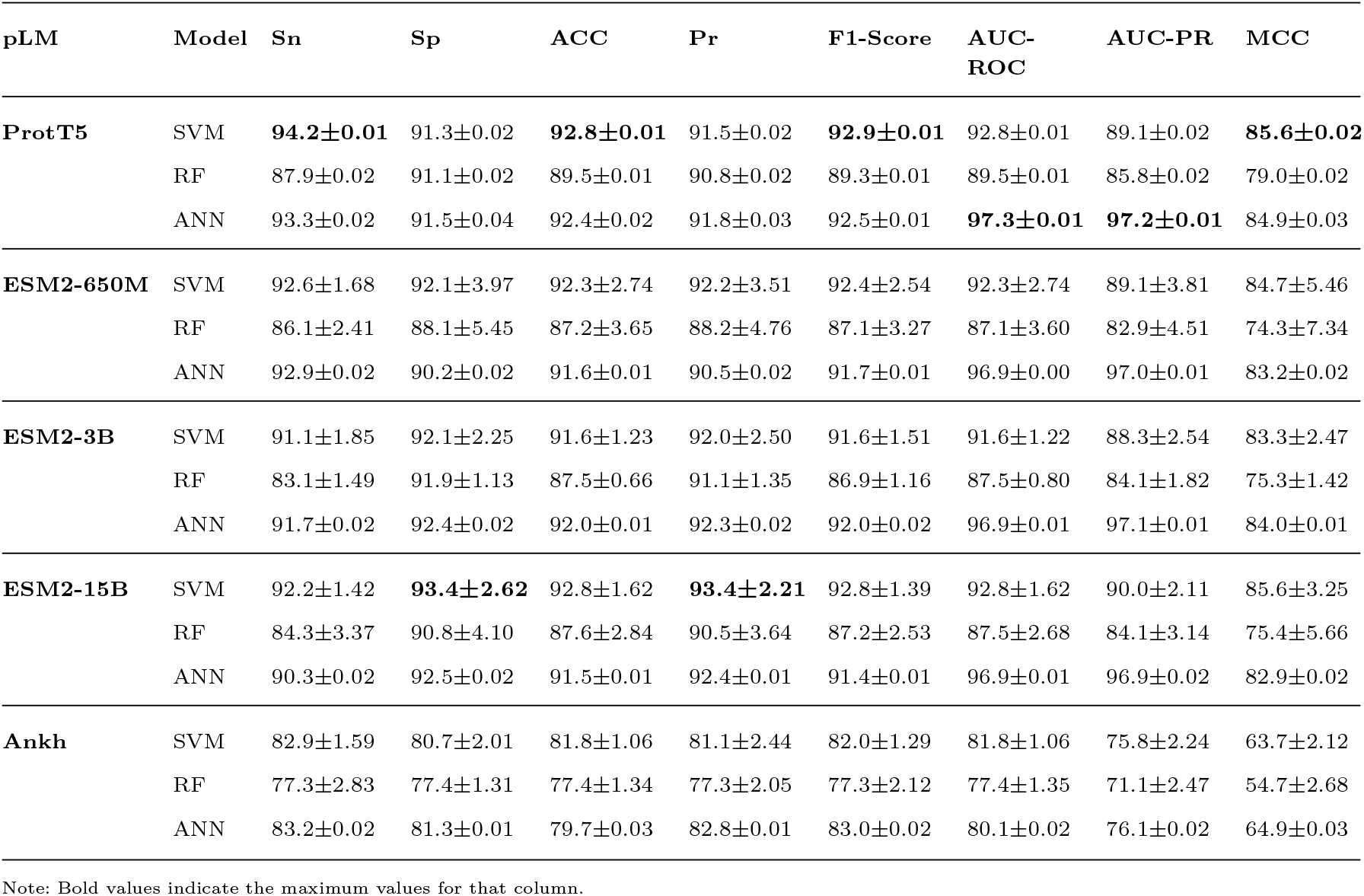
The comparison of 5-fold cross-validation results based on embeddings from ProtT5, Ankh, ESM2-650M, ESM2-3B, and ESM2-15B.

| pLM | Model | Sn | Sp | ACC | Pr | F1-Score | AUC-ROC | AUC-PR | MCC |
| --- | --- | --- | --- | --- | --- | --- | --- | --- | --- |
| **ProtT5** | SVM | **94.2±0.01** | 91.3±0.02 | **92.8±0.01** | 91.5±0.02 | **92.9±0.01** | 92.8±0.01 | 89.1±0.02 | **85.6±0.02** |
|  | RF | 87.9±0.02 | 91.1±0.02 | 89.5±0.01 | 90.8±0.02 | 89.3±0.01 | 89.5±0.01 | 85.8±0.02 | 79.0±0.02 |
|  | ANN | 93.3±0.02 | 91.5±0.04 | 92.4±0.02 | 91.8±0.03 | 92.5±0.01 | **97.3±0.01** | **97.2±0.01** | 84.9±0.03 |
| **ESM2-650M** | SVM | 92.6±1.68 | 92.1±3.97 | 92.3±2.74 | 92.2±3.51 | 92.4±2.54 | 92.3±2.74 | 89.1±3.81 | 84.7±5.46 |
|  | RF | 86.1±2.41 | 88.1±5.45 | 87.2±3.65 | 88.2±4.76 | 87.1±3.27 | 87.1±3.60 | 82.9±4.51 | 74.3±7.34 |
|  | ANN | 92.9±0.02 | 90.2±0.02 | 91.6±0.01 | 90.5±0.02 | 91.7±0.01 | 96.9±0.00 | 97.0±0.01 | 83.2±0.02 |
| **ESM2-3B** | SVM | 91.1±1.85 | 92.1±2.25 | 91.6±1.23 | 92.0±2.50 | 91.6±1.51 | 91.6±1.22 | 88.3±2.54 | 83.3±2.47 |
|  | RF | 83.1±1.49 | 91.9±1.13 | 87.5±0.66 | 91.1±1.35 | 86.9±1.16 | 87.5±0.80 | 84.1±1.82 | 75.3±1.42 |
|  | ANN | 91.7±0.02 | 92.4±0.02 | 92.0±0.01 | 92.3±0.02 | 92.0±0.02 | 96.9±0.01 | 97.1±0.01 | 84.0±0.01 |
| **ESM2-15B** | SVM | 92.2±1.42 | **93.4±2.62** | 92.8±1.62 | **93.4±2.21** | 92.8±1.39 | 92.8±1.62 | 90.0±2.11 | 85.6±3.25 |
|  | RF | 84.3±3.37 | 90.8±4.10 | 87.6±2.84 | 90.5±3.64 | 87.2±2.53 | 87.5±2.68 | 84.1±3.14 | 75.4±5.66 |
|  | ANN | 90.3±0.02 | 92.5±0.02 | 91.5±0.01 | 92.4±0.01 | 91.4±0.01 | 96.9±0.01 | 96.9±0.02 | 82.9±0.02 |
| **Ankh** | SVM | 82.9±1.59 | 80.7±2.01 | 81.8±1.06 | 81.1±2.44 | 82.0±1.29 | 81.8±1.06 | 75.8±2.24 | 63.7±2.12 |
|  | RF | 77.3±2.83 | 77.4±1.31 | 77.4±1.34 | 77.3±2.05 | 77.3±2.12 | 77.4±1.35 | 71.1±2.47 | 54.7±2.68 |
|  | ANN | 83.2±0.02 | 81.3±0.01 | 79.7±0.03 | 82.8±0.01 | 83.0±0.02 | 80.1±0.02 | 76.1±0.02 | 64.9±0.03 |

Note: Bold values indicate the maximum values for that column.

Furthermore, consistent with the findings of Jahn et al. [7], we also observed that the choice of pLM is not critical. ESM yielded comparable results to ProtT5, with ProtT5 showing slightly better performance, though the difference was not statistically significant. However, the dimensionality of the features might be another criterion for the selection of pLM. In Figure 2, we can observe that ProtT5 features are exhibiting higher classification performance in terms of most of the measured features. Although ESM 15B has on-par performance, because of the additional computational cost of using ESM 15B over ProtT5, we selected ProtT5 representations as the features to develop our final model.

**Fig. 2.**
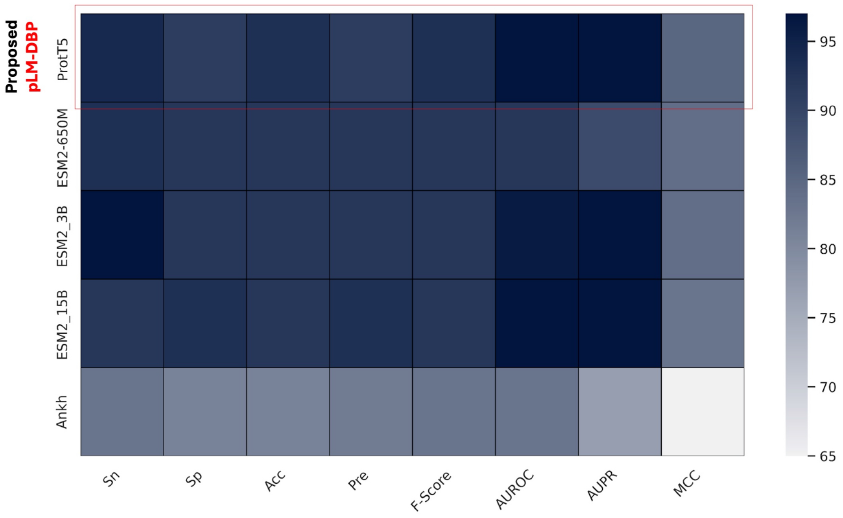
Comparison of the highest performance achieved from Ankh, ESM2 3B, ESM2 3B, ESM2 650M, and ProtT5 based on Table 4.

Note: Bold values indicate the maximum values for that column.

### Analysis of features

We used UMAP (Uniform Manifold Approximation and

Projection) [12] to visualize high-dimensional features in a lower-dimensional space. Specifically, we visualized the 2D UMAP projections of both the original ProtT5 representations and the refined pLM-DBPs representations in a scatter plot (See Figure 3), highlighting the distinction between DBPs (represented in blue) and NDBPs (represented in orange). As shown in Figure 3-a, the ProtT5 features alone exhibit clear separation between DBP and NDBP clusters. After training the pLM-DBPs model using ProtT5 features, we observe an even clearer distinction between DNA-binding proteins (DBPs) and non-DNA-binding proteins (NDBPs) in the resulting scatter plot. Even though this is merely a visualization in a lower-dimensional space, the observed separation highlights the efficiency of our proposed approach in uncovering insights into the underlying biological characteristics of DBPs and NDBPs.

**Fig. 3.**
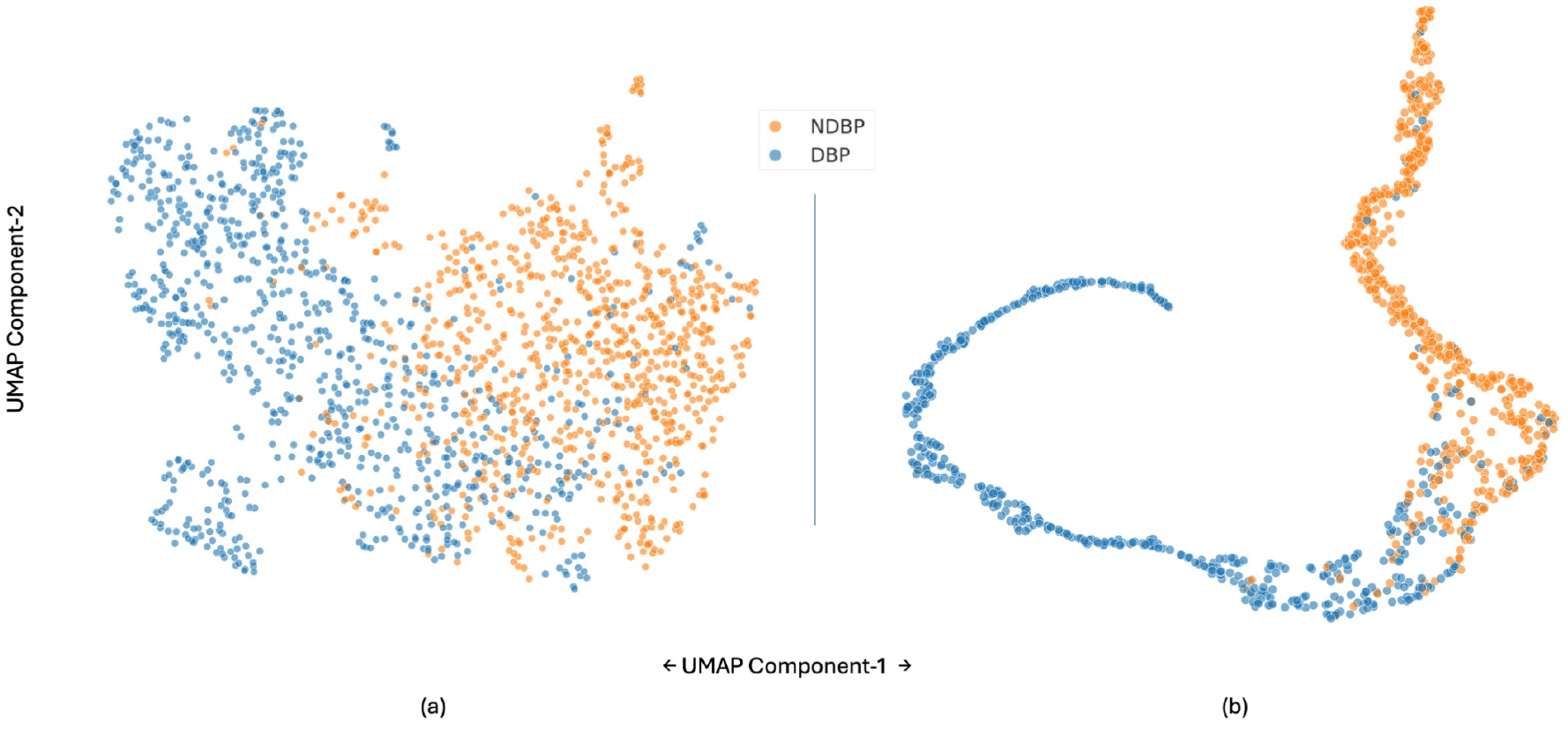
Embedding evolution plot, illustrating 2D UMAP projections of learned features. (a) Visualization of ProtT5 embeddings and (b) embeddings from the penultimate layer of pLM-DBPs. The plots depict how the feature representations evolve, highlighting class separation and clustering patterns as the model refines its understanding of DNA-binding (DBP) and non-DNA-binding proteins (NDBP).

### Comparison with existing approach using the independent test set

We compared the results of our proposed model with existing tools DNAbinder [8], DPP_PseAAC [22], IDRBP_MMC [29], IDRBP-PPCT [25], PDBP-fusion [9], DeepDRBP [28], StackDPPred [13], and PlDBPred [19] on the independent test set described in Table 5. Most of the comparison results are adopted from PlDBPred [19].

Our comparative analysis evaluated the proposed model against existing approaches using several key metrics: sensitivity, specificity, accuracy, precision, F-Score, and MCC. Although AUROC and AUPR are not included in Table (5) as some of the existing tools that we are comparing against have not reported these metrics. However, pLM-DBPs achieved impressive AUROC and AUPR values of 96.7 and 97.3, respectively on the independent test set. Furthermore, Figure 4—a radar plot illustrating the performance across the metrics from Table 5—demonstrates that pLM-DBPs outperform other tools comprehensively, excelling in all evaluated metrics, including accuracy, specificity, precision, MCC, and sensitivity.

**Table 5.**
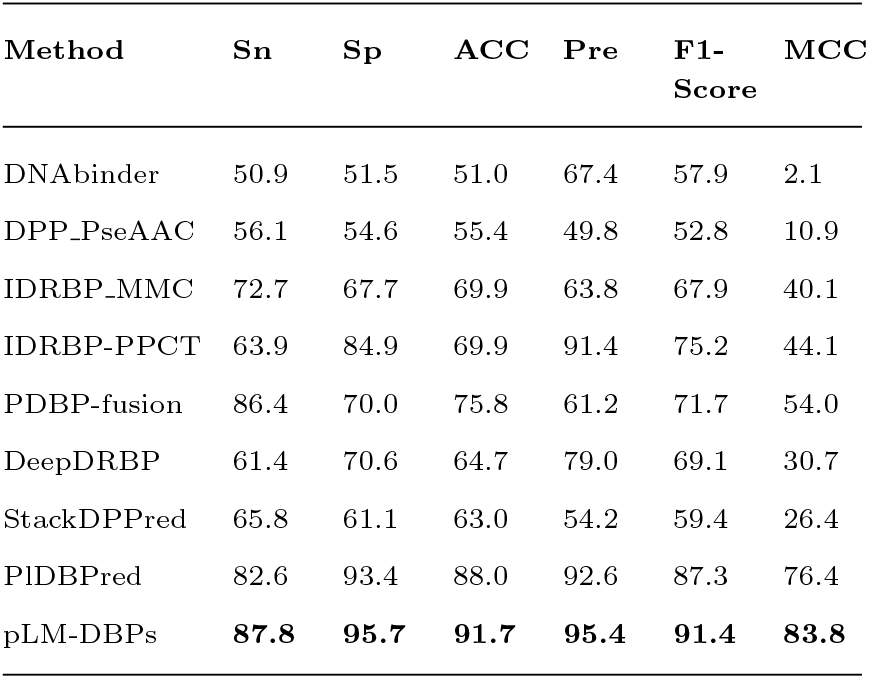
Comparison of performance metrics between pLM-DBPs and existing tools on independent Test Set.

**Fig. 4.**
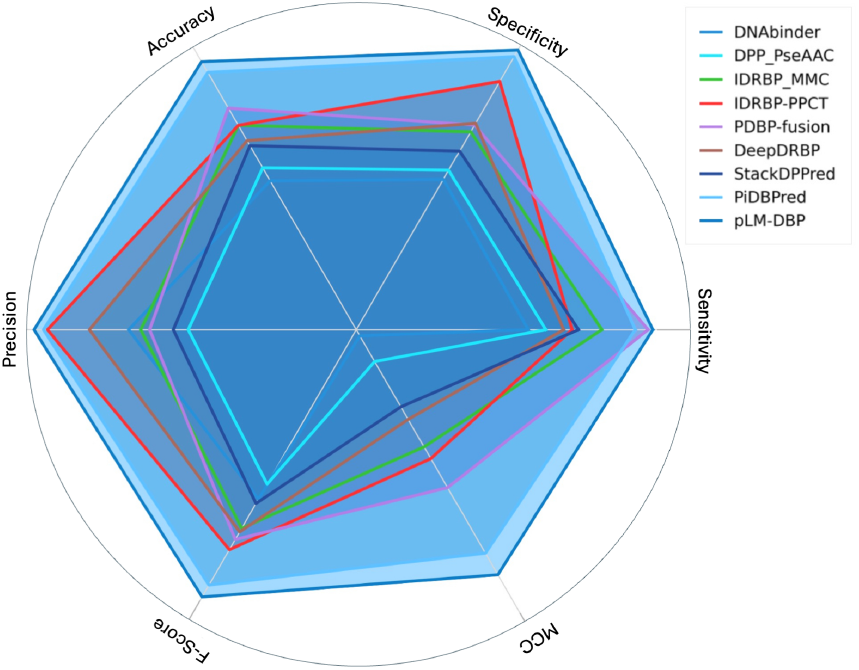
Radar plot comparing the performance of pLM-DBPs with existing methods using key metrics: sensitivity (Sn), specificity (Sp), accuracy (ACC), precision (Pre), F-Score, and Matthews Correlation Coefficient (MCC). The plot demonstrates the superior performance of pLM-DBPs across all metrics, particularly excelling in accuracy, specificity, precision, and MCC, when compared to other approaches such as DNAbinder, DPP_PseAAC, IDRBP_MMC, IDRBP-PPCT, PDBP-fusion, DeepDRBP, StackDPPred, and PiDBPred.

### Evaluation of methodology on the multiclass DNA/RNA/SSB dataset

To further assess the effectiveness of our proposed methodology

—ProtT5 features coupled with an ANN classifier—we applied it to the multiclass dataset of DNA-binding as recently analyzed by Wu and Guo (Wu and Guo 2024). This dataset consists of four classes: RNA binding proteins (RBP), single-stranded DNA binding proteins (SSB), double-stranded DNA binding proteins (DSB), and non-NABPs (proteins without DNA/RNA binding properties). We followed their approach by splitting the dataset into 81% training, 9% validation, and 10% test sets for a fair comparison.

When evaluated on the test set, our method achieved accuracies of 91.46% for RBP, 93.35% for DSB, and 85.71% for SSB. These results demonstrate a substantial improvement over the accuracies reported by Wu and Guo (Wu and Guo 2024) (RBP: 70.3%, DSB: 79.6%, and SSB: 39.4%). Specifically, our approach improved the classification accuracy for RBP by 30.1%, DSB by 17.27%, and SSB by 117.58%. Similar improvements were also observed in the validation set, reinforcing the robustness of our method. Furthermore, the 2D UMAP features visualization in Figure 5 demonstrates well-separated clusters, emphasizing the effectiveness of the proposed methodology in a multiclass classification objective. The distinct clusters correspond to the different classes: RBP (Green), DSB (Blue), SSB (Red), and NABP (Orange). The features used for this visualization were extracted from the penultimate layer of the proposed model trained using a multiclass dataset and subsequently reduced to two dimensions using UMAP. These results suggest that our methodology is highly effective for plant-based DNA-binding predictions and can also generalize well to other DNA-binding and RNA-binding tasks, including multiclass classification problems.

**Fig. 5.**
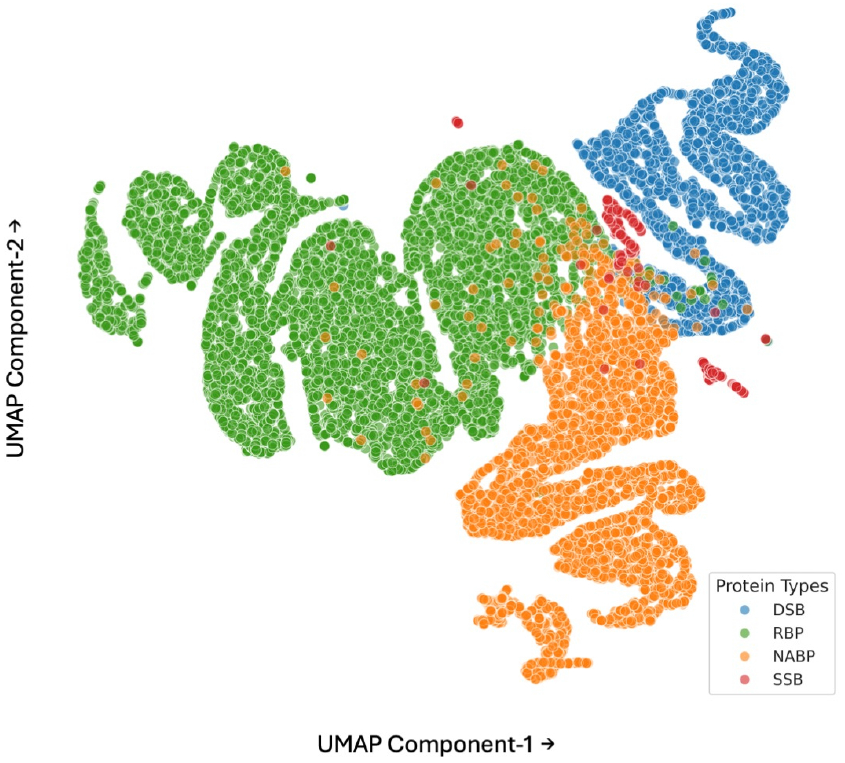
2D UMAP features visualization of the penultimate layer features, showing well-separated clusters for RBP (Green), DSB (Blue), SSB (Red), and NABP (Orange) classes, demonstrating the model’s effectiveness in multiclass classification.

### Conclusion and future work

In this work, we developed pLM-DBPs, a robust prediction model for DNA-binding proteins in plants. After analyzing features from prominent protein language models such as ESM2, ProtT5, and Ankh, we designed an ANN-based predictor trained on ProtT5 embeddings that efficiently identifies plant DNA-binding proteins with high accuracy. We experimented with multiple machine learning models and a broad range of hyperparameters, ultimately we developed a simple yet highly effective ANN model for plant DBP prediction. We compared the performance of pLM-DBPs with existing DBP prediction tools and also validated our approach using other available datasets for DNA-binding prediction tasks where we consistently observed the superior performance of our model.

This work opens several avenues for future research for DNA-binding protein prediction tasks. One promising direction is the exploration of other available protein language models that are not explored in this study, such as ESM3, to potentially enhance prediction accuracy. Secondly, leveraging protein 3D-structure-based models could provide deeper insights, particularly in capturing the 3D structural information on DNA-binding interactions. Another valuable area for further investigation is the integration of multiple protein language models, as described by Pokharel et al. [18], that can potentially offer a combined strength of multiple models to enhance the predictive capabilities. These directions present exciting opportunities to build upon the findings of this study and drive further progress in the field.

## Author Contributions Statement

D.K. and S.P. conceived the experiment(s), S.P., K.B. conducted the experiment(s). K.B. and P.P. performed feature extraction. K.B. and S.P. wrote the manuscript. D.K., S.P., P.P. revised the manuscript. D.K. oversaw the project.

## Acknowledgments

We thank Dr. Stefan Schulze for the helpful discussion during the development of this project. We acknowledge the use of the DeepBlizzard HPC Cluster at Michigan Tech. We also acknowledge the authors of PlDBPred for making the dataset available.

## Supplementary Materials

Supplementary information is provided online which can also be found at https://github.com/suresh-pokharel/pLM-DBPs.

## Data Availability

Data, codes, and other resources are publicly available at https://github.com/suresh-pokharel/pLM-DBPs. Information about the web server will be provided through the same GitHub repository.

## Footnotes

GitHub: https://github.com/suresh-pokharel/pLM-DBPs

