## Supplementary Information for "pLM-DBPs: Enhanced DNA-Binding Protein Prediction in Plants Using Embeddings From Protein Language Models"

**Table T1: The comparison of 5-fold cross-validation results based on attention pooling features from ProtT5, Ankh, ESM2-650M, ESM2-3B, and ESM2-15B**

| pLM | Model | **Sn** | **Sp** | **ACC** | **Pr** | **F-Score** | **AUC-ROC** | **AUC-PR** | **MCC** |
|---|---|---|---|---|---|---|---|---|---|
| ProtT5 | SVM | **93.0±3.04** | **93.2±1.05** | **93.2±1.78** | **93.2±1.07** | **93.1±1.96** | 93.1±1.78 | 90.2±2.13 | **84.3±3.48** |
| | RF | 88.3±4.15 | 90.7±1.26 | 89.6±2.27 | 90.5±1.24 | 89.4±2.66 | 89.5±2.24 | 85.8±2.81 | 79.1±4.34 |
| | ANN | 87.6±0.07 | 92.6±0.03 | 90.1±0.02 | 92.4±0.03 | 89.7±0.03 | 96.0±0.01 | 96.3±0.01 | 80.5±0.04 |
| ESM2-650M | SVM | 90.3±1.64 | 92.6±3.59 | 91.4±2.30 | 92.5±3.43 | 91.3±2.12 | 91.4±2.31 | 88.4±3.39 | 82.9±4.62 |
| | RF | 82.4±2.47 | 93.2±2.74 | 87.8±1.48 | 92.4±2.73 | 87.1±1.79 | 87.8±1.59 | 84.9±2.66 | 76.1±3.13 |
| ESM2-3B | SVM | 90.2±3.29 | 92.3±2.16 | 91.2±1.97 | 92.0±2.70 | 91.1±2.16 | 91.3±1.94 | 87.9±3.02 | 82.5±3.85 |
| | RF | 83.4±2.26 | 91.9±2.08 | 87.6±0.96 | 91.1±2.60 | 87.0±0.70 | 87.7±0.76 | 84.3±1.82 | 75.5±1.69 |
| | ANN | 91.6±0.01 | 92.9±0.02 | 92.2±0.01 | 92.9±0.01 | 92.2±0.01 | **96.9±0.01** | **96.9±0.01** | 84.5±0.02 |
| ESM2-15B | SVM | 90.8±2.14 | 92.6±3.16 | 91.7±2.31 | 92.5±3.31 | 91.6±2.36 | 91.7±2.28 | 88.6±3.60 | 83.4±4.63 |
| | RF | 85.4±2.88 | 90.3±1.65 | 87.8±1.08 | 89.7±2.08 | 87.5±1.43 | 87.8±1.09 | 83.9±2.03 | 75.7±2.10 |
| Ankh | SVM | 80.5±0.04 | 79.8±0.04 | 80.2±0.03 | 79.9±0.05 | 80.1±0.03 | 80.2±0.03 | 74.1±0.05 | 60.3±0.05 |
| | RF | 76.2±0.05 | 78.8±0.03 | 77.6±0.04 | 78.1±0.05 | 77.1±0.05 | 77.5±0.04 | 71.5±0.06 | 55.0±0.07 |

**Table T2: Hyperparameters used to train the final ANN model (pLMDBPs). These hyperparameters were selected after experimenting with several models with a wide range of configurations using grid search and KerasTuner.**

| | |
|---|---|
| No. of Hidden Layers | 4 |
| No. of Neurons in Hidden Layers | 512, 512, 128, 16 |
| Dropout Rates at Each Hidden Layer | 0.3, 0.3, 0.4, 0.4 |
| Input/output Size | 1024 / 1 |
| Loss Function | BinaryCrossEntropy() |
| Activation Functions in Intermediate Layers | ReLu |
| Activation Function in the output layer | Sigmoid |
| Training Optimizer | Adam(learning_rate=0.00001) |
| No. of Epochs | 86 |
| Callback Method | Early Stopping |
| Trainable/ optimizer Parameters: | 855,201 |

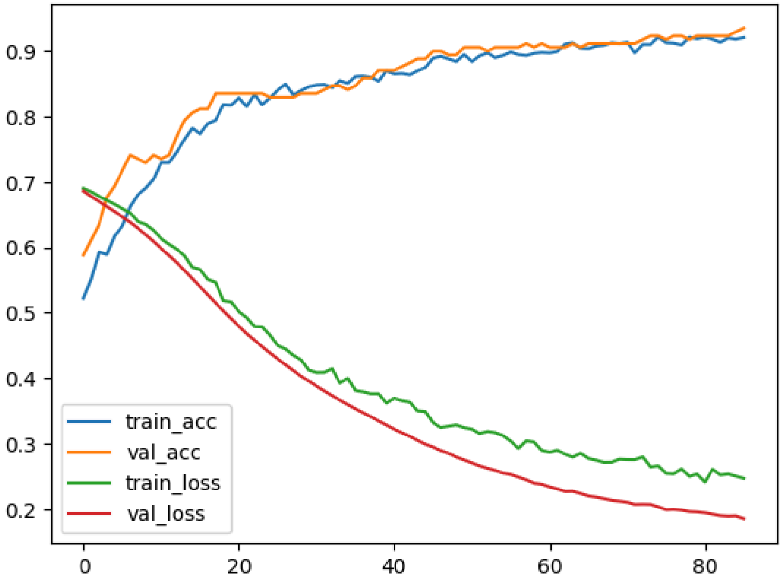


**Figure S1:** Training statistics of the pLMDBPs presenting accuracy and loss curve for training and validation. We can observe a smooth decrease in loss and a consistent increase in accuracy, and close alignment between the training and validation accuracy/loss indicated that the model has learned effectively without overfitting or underfitting during the training.
